# A novel in vitro 3D cancer model based on modular tissue engineering approach

**DOI:** 10.1101/2025.02.25.640239

**Authors:** N Daneshvar Baghbadorani, M Bosso, R Greene, T Dzikowski, M Johannson, A Gagnon, MD Chamberlain

## Abstract

An emerging tool to better recapitulate the complexity of tumor biology in vitro is 3D culture models. Here, we describe a free-floating collagen-based hydrogel system with embedded cancer cells, called microtissues. The microtissues are based on the well-established modular tissue engineering method. They mimic the natural development of the tumor microenvironment, with features such as hypoxia and treatment resistance. To demonstrate the utility of microtissues as a 3D tumor model system, triple negative breast cancer cells were cultured using this method and were shown to maintain cell viability and proliferation with minimal cell death, along with mimicking natural emergence of tumor properties such as, a hypoxic core. Furthermore, by screening the model with commonly used anti-breast cancer chemotherapeutics, we observed drug resistance to concentrations which are largely in accordance with the used doses in the clinics. Therefore, our model offers the opportunity to naturally reproduce fundamental features of a tumor in vitro, leading to emergence of a similar cell reprogramming which is responsible for clinical drug resistance.

## Introduction

The majority of novel drug candidates, especially anti-cancer therapies, fail to establish clinical therapeutic efficacy. This issue is partly due to the inaccurate preliminary assessments of novel drug candidates by techniques and models which are not able to resemble the complexity of human pathologies^1–3^. In the case of cancer, despite the historical success of 2D-cultured cancer cell lines, which has characterized the main part of our current knowledge of cancer biology, this simple in vitro model is associated with challenging drawbacks^4^. One of the biggest challenges is that tumors are a heterogenous population of various cell types, including cancer, stromal and immune cells supported by a three-dimensional (3D) network of macromolecules, the extracellular matrix (ECM)^5^. The 3D geometry of the tumor allows interaction between cells and their surrounding microenvironment, and establishment of nutrient and oxygen gradients^6^. Additionally, this 3D microenvironment is the major determinant of tumor cell behaviour, promoter of cancer progression, therapy resistance and disease relapse^7–9^. However, in 2D cancer models, a homogeneous group of cancer cells is cultured as a monolayer of adherent cells in a dish with unlimited access to oxygen and nutrients, and in the absence of cancer associated stromal cells and ECM that make up the tumor microenvironment (TME)^4^. On the other hand, in vivo studies as the other major part of preliminary drug studies are limited by ethical, time, and cost-effectiveness issues, although they provide us with a partly functional TME of murine origin. The murine TME is similar to the human one, but not identical^10,11^ and immunocompromised mice are often used to grow the tumors. Consequently, the development of 3D in vitro models able to mimic the main features of TME, including the extracellular matrix (ECM), oxygen and nutrient gradients, hypoxia, and immune cell surveillance has been considered as a solution to overcome limitations of 2D in vitro models and animal studies.

Thanks to recent advances, tissue-engineered 3D models are being developed to characterize a tumor microenvironment to better reproduce the biological behaviour of tumor cells. These models have different levels of complexity starting from simple tumor spheroids^12^ and moving towards more advanced scaffold-based models^13^, microfluidics^14,15^, and organoids^16,17^. Among the variety of introduced scaffolds for 3D cell culture, collagen-based hydrogels have been shown by several studies^18–21^ as a promising biocompatible scaffold to recapitulate the microenvironment of a solid tumor. Collagen is one of the main components of natural extracellular matrix and the amount of collagen is increased compared to normal tissue in several tumor types^22^. Our study is focused on the implementation of modular tissue engineering designs to the field of cancer research to overcome some of the limitations of conventional 3D-cell culture. Although this technique has been initially developed for tissue-engineering and transplantation research^23–26^, we believe it has several advantages to offer in the field of 3D tumor cell culture. The model system stemming from this approach is a free-floating, cylindrical collagen hydrogel which can be embedded with cancer cells and other cell types to mimic the TME. These collagen hydrogel constructs are called microtissues. The microtissues are easy to use and manipulate similar to 2D cell culture, can be used in any standard molecular or cellular assay and are cost effective to make with no need for expensive manufacturing equipment. In this paper, we describe the features of the in vitro tumor model which was developed by the modular tissue engineering approach. This basic model contains only cancer cells embedded in the collagen hydrogel. Using this model, we cultured the triple negative breast cancer cell lines, HCC1806 and MDA-MB-231, and assessed the microtissues for viability and proliferation, morphological alterations, emergence of hypoxia, and resistance to chemotherapeutic agents.

## Materials and Methods

### Drugs and Reagents

Stock solutions of Vincristine (Focus Biomolecules, CAT# 10-2570) and Vinblastine (LC Laboratories, CAT# V-7300) were prepared in water. Stock solution of Paclitaxel (Cayman Chemical, CAT# 10461), Doxorubicin (APExBIO, CAT# A3966) were prepared in DMSO (Sigma-Aldrich). All stock solutions were kept at -80 °C.

### Cell culture

Experiments were conducted using two human triple negative breast cancer cell lines and both cell lines were cultured using the specifications outlined by ATCC. The HCC1806 cell line was cultured in RPMI 1640 (Gibco) medium supplemented with 10% fetal bovine serum (Gibco) and 1% Penicillin/Streptomycin (Invitrogen) and was maintained at humidified atmosphere at 37 °C and 5% CO_2_. The MDA-MB-231 cell line was cultured in L-15 Leibovitz (Gibco) medium supplemented with 10% fetal bovine serum (Gibco) and 1% Penicillin/Streptomycin (Invitrogen) and was maintained in a non-CO_2_ incubator at 37 °C.

### Fabrication of microtissues

Microtissues of HCC1806 or MDA-MB-231 were fabricated similar to previous methods^27,28^ with a few modifications. Briefly, Bovine type I Collagen (3 mg/mL, PureCol™) was mixed with 10X minimum essential media (MEM, Gibco) stock solution to a 1X MEM solution, and neutralized by 1M sodium bicarbonate solution to achieve pH 7.4 and kept on ice to prevent gelation. The cells were added at a density of 2 million cells/mL of collagen, the mixture was loaded into sterile polyethylene tubing (PE90, BD Bioscience), and incubated in a humidified atmosphere at 37 °C and 5% CO_2_ for 1 hour to gel. The tubing was then cut using a specialized 3D-printed cutting plate (Fig. S1). The cutting plate holds the tubing and has cutting tracks that guide the razor blade through designated slots which were 2 mm apart. The cut pieces were collected in media in a 50 ml tube, and vortexed to remove the tubing from the microtissues. The microtissues where then maintained in the conditions required by the cell line as outlined by ATCC.

### Histology

Microtissues were washed with PBS and fixed in 10% Neutral Buffered Formalin (NBF) overnight. The microtissues were then embedded in 1.5% agarose using a cryo-mould. The agarose blocks were fixed in 10% NBF for 24 hours, washed in 70% ethanol twice, and left in 70% ethanol until further processing. After processing, slides where either stained for H&E or immunohistochemistry (IHC). For IHC, heat-induced epitope retrieval was performed by microwave treatment in sodium citrate buffer (10 mM Sodium citrate, 0.05% Tween 20, pH 6.0), followed by washing and blocking of endogenous peroxidases with 3% hydrogen peroxide. Tissue sections were then blocked with 3% BSA and incubated with Ki67 (1:1000; Thermofisher, Cat# PA5-19462), active Caspase-3 (1:200, Thermofisher, Cat# MA5-32015), or processed with DeadEnd™ Colorimetric TUNEL System (Promega, Cat# G7360). After washing, slides were incubated with an HRP-labeled secondary antibody, washed again, and developed with DAB chromogen (DAKO EnVision™+). Slides were scanned using the Aperio virtual microscope (CS Scanscope).

### Viability assays

The viability of microtissues was assessed at day 1, 3, 5, 7, 10, 14 and 21. At each day, three microtissues were transferred to a well of a non-tissue culture treated 96-well plate in 100 μL of media. At least three wells were considered for each experiment. After seeding, the viability was assessed by the CellTiter-Glo® 3D (Promega) kit according to its manufacturer`s protocol. Each well received 100 μL of reagent followed by vigorous shaking for 5 mins, then microtissues were transferred to an opaque 96-well plate and incubated at room temperature for 25 mins. Luminescence was read by the Varioskan LUX multimode microplate reader.

### Live-Dead staining assay

Microtissues were transferred to a chambered coverglass-bottomed slide (Lab-Tek™), washed with PBS, and incubated with Calcein acetoxymethyl ester (1:2000), and Propidium iodide (1:2000) for 30 minutes according to the live/dead kit instructions (Thermofisher, Cat# L32250). The assay was performed at day 0, 3, 5, 10,14 and 21. Following incubation, microtissues were imaged by confocal microscopy (Zeiss LSM700).

### Drug screening assay

Microtissues were cultured for 5 days and then three microtissues in 100 μL of culture media were transferred into the wells of a non-tissue culture treated 96-well plate. In the case of 2D-cultured cells, 2000 cells in 100 μL per well of a tissue-culture treated 96-well plate were used. A 7-point serial dilution of the drug was prepared in media at double the final concentration and 100 μL of the treatment were added to each well in triplicate. Control treatments were cell media only, and vehicle control, which was the drug vehicle (DMSO or water) at the same concentration as the highest drug concentration. The treated microtissues were incubated at 37 °C and 5% CO_2_ for 4 days. The effect of the drug was assessed by CellTiter-Glo® 3D (Promega) as per manufacturer’s instructions as outlined above. The assay results were normalized to the average of the media control wells.

### Assessment of Hypoxia

Presence of hypoxia in microtissues was evaluated by HypoxiTRAK^TM^ (Biostatus), a molecular probe which converts to a fluorescent metabolite in hypoxic cells and irreversibly accumulates. Microtissues for this assay were cultured from day 0 in phenol red-free media to prevent possible false positive fluorescent signals of phenol red. Starting at day 5, the microtissues were cultured with 50 nM HypoxiTRAK^TM^ in phenol red-free media. Nuclei were stained with Hoechst 33,342. Hypoxia was assessed at day 7, 10, 14 and 21 by confocal microscopy (Zeiss LSM700).

### Doxorubicin diffusion assay

Microtissue-cultured HCC1806 cells at day 5 were treated with 2.5 μM doxorubicin, and nuclei were stained with Hoechst 33,342. After 30 minutes incubation in a humidified atmosphere at 37 °C and 5% CO2, live cell imaging was done for 12 hours by Confocal microscopy (Zeiss LSM700) with an image of the middle section of the microtissue taken every 30 minutes.

### Statistical analysis

Data was analysed by GraphPad prism software 9. P-values were determined by two-way ANOVA followed by Tukey’s multiple comparisons (** p< 0.01, **** p < 0.0001).

## Results

### Cellular architecture of the microtissues

The microtissues start out 2 mm long and 0.8 mm wide in size but shrink as the cells remodel the tissue (Fig. S2). This shrinkage of the microtissue is caused by the cells as they proliferate and form structures within the tissue. As the cells engage with the collagen, they put contractile forces on the hydrogel. The shrinkage occurs because the microtissues are free-floating and are not attached to the walls or bottom of the container. This attachment to the container that adds structural rigidity to the hydrogel limits the contraction of the ECM hydrogel. This has been seen with other 3D culture systems^29^. The change in size of the microtissue is cell type dependent and the size of the microtissue stabilizes within the first 5 days of growth. The histological examination of microtissue-cultured HCC1806 cells (Fig. 1) over a 21-day period showed that by day 10, clusters of cells homogenously populated the collagen hydrogel and completely filled the structure. However, at later time points, cell clusters and their organized distribution were lost, and cells started to concentrate on the edges of the microtissue. Consequently, a hollow space started to reappear within the core, suggesting the migration of cells towards the edges of the structure. Similar results were seen with MDA-MB-231 cells however they typically formed only small clusters of cells compare to the HCC1806 (Fig. 1). This hollowing out of the microtissues could indicate the establishment of an oxygen and nutrient gradient, and possibly a hypoxic core which forces the cells to migrate toward the edges similar to an actual tumor.

**Figure 1.**
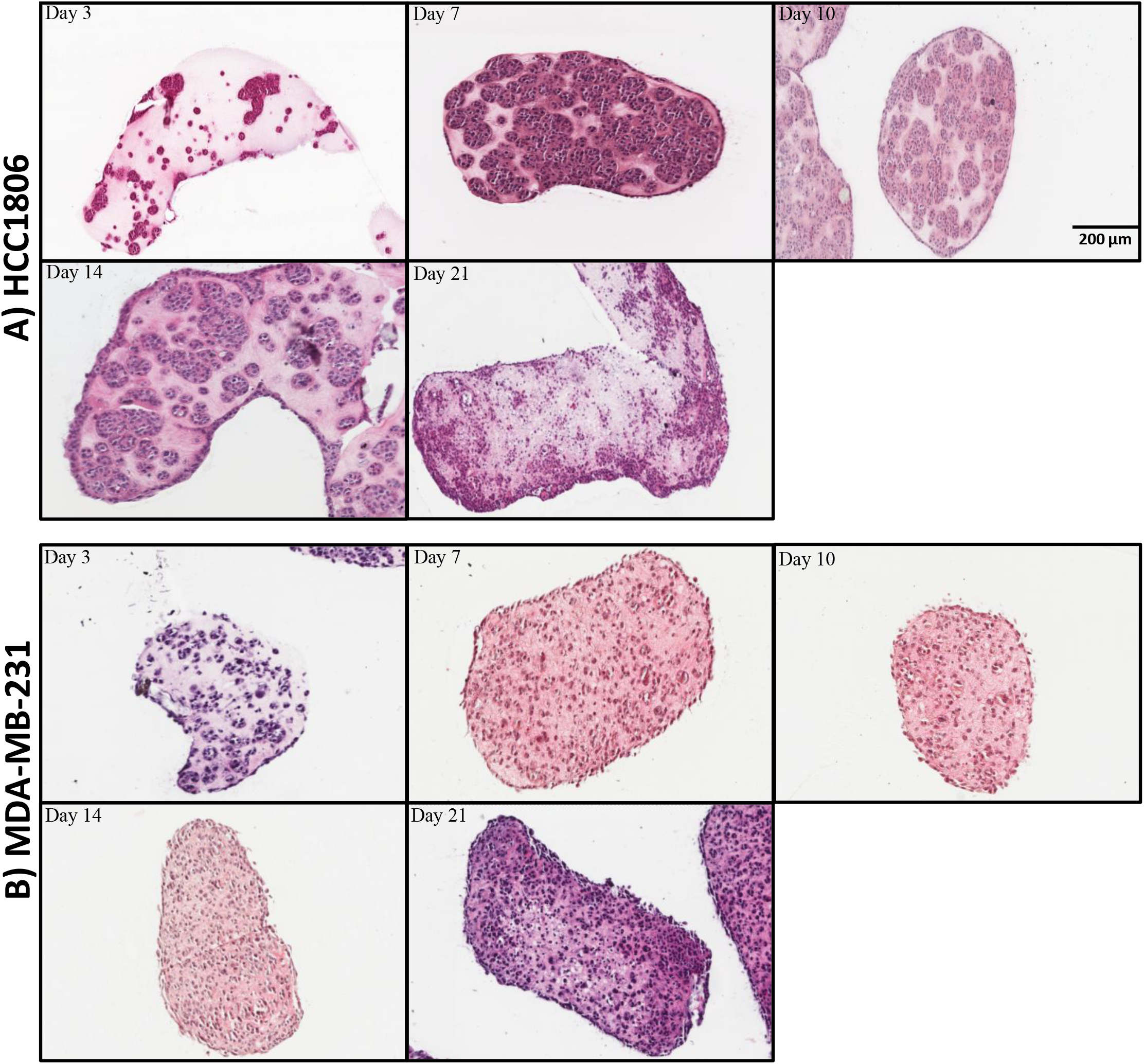
Morphology of Microtissues. H&E staining of microtissue-cultured HCC1806 (A) and MDA-MB-231 (B) cells at day 3, 7, 10, 14 and 21 (N=1). A) As the HCC1806 microtissues age, the cells proliferate and grow in clusters. The clusters are highest in number at day 10, but cell clustering diminishes at later time points, and by day 21 most of the cells are found along the edges of the structure.

### Microtissue-cultured cancer cells retain their viability and proliferative capacity

It was essential to investigate the viability of cells in microtissues to demonstrate the capacity of our model in keeping cells alive. The viability of two different triple negative breast cancer cell lines, HCC1806 and MDA-MB-231 cells, was assessed over a 21-day period (Fig. 2). A live/dead staining assay was initially performed to evaluate viability of microtissue-cultured cells (Fig. 2A,B). As seen with other cell types grown in modular tissue engineered constructs^30,31^, both tested cell lines maintained good cell viability after the formation of the microtissues. Additionally, this observation was confirmed by an ATP release assay to measure cell viability. As shown in figure 2C, the HCC1806 cells proliferate over time as there is a continuous increase in ATP levels over the first 10 days and then the ATP levels plateaus. In the case of MDA-MB-231, a certain level of ATP release was maintained over the 21-day growth period. The significant difference between the levels of released ATP between the two cell lines could be due to their different proliferation rates. Overall, cells maintain their proliferation and viability in this model, but the stabilization of the cell number is cell line dependent. This stabilization could be due to decreases in proliferation, a steady state turnover of cells, or continued proliferation of the cells with a decrease in cell metabolism as the level of ATP in the cells is used as a proxy for cell viability.

**Figure 2.**
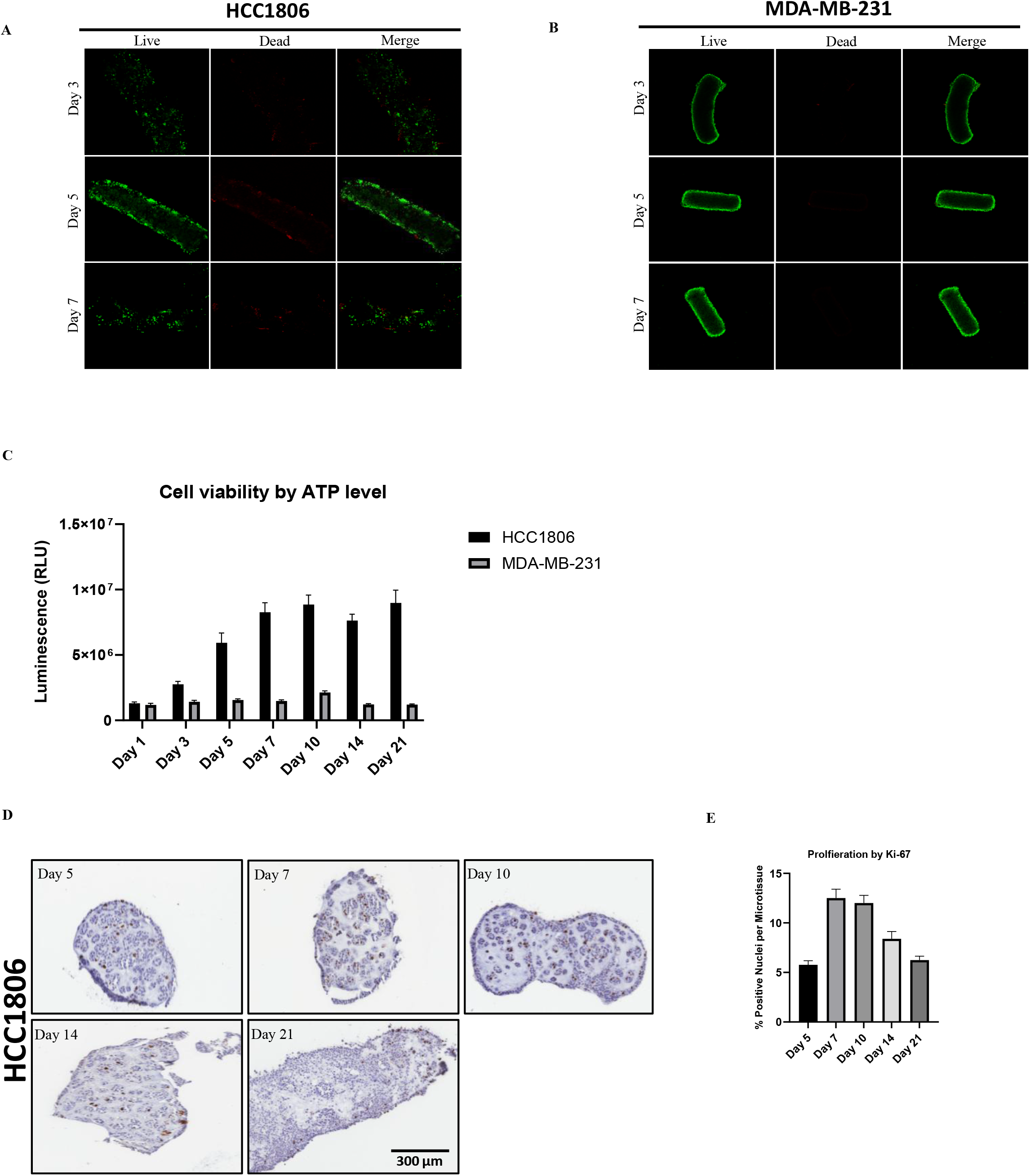
Assessment of viability and proliferation. Representative images of live/dead staining of the 3D microtissue-cultured HCC1806 (A) and MDA-MB-231 (B) cells at day 3, 5, 7 taken by confocal microscope (N=2). The green staining is Calcein AM for live cells and the red staining is ethidium homodimer-1 for dead cells. (5X) C) The viability of HCC1806 and MDA-MB-231 cells cultured in microtissues were measured based on ATP release using CellTiter-Glo 3D during a 21-day growth period. Data are reported as mean ± SEM of 3 experimental replicates. D) Representative images of Ki67 staining of 3D microtissue-cultured HCC1806 cells (5X). Ki67 as the prominent marker of proliferating cells was stained using Immunohistochemistry (IHC) (N=3). Brown nuclei represent proliferative cells. E) Quantification of percentage of KI-67 positively stained cells. Each replicate involved counting the total number of KI-67 positive cells normalized by the total number of cells of one microtissue. Data represent mean ± SEM of at least three experimental replicates.

The proliferation of the cells within the HCC1806 microtissue was also confirmed by Ki-67, a marker of cell proliferation (Fig. 2D,E). Ki-67 is a lagging indicator of cell proliferation, and the number of Ki-67^+^ cells peaks by day 7 to 10 and then decreases between day 10 and 21 (Fig. 2E), which supports the observed variations of ATP levels in HCC1806. When the distribution of the Ki-67^+^ cells in the microtissues was studied the ki-67^+^ cells were roughly evenly distributed across the tissue for the first 14 days showing that the proliferating cells where both in the centre and edge of the tissue. However, by day 21 the ki-67^+^ cells where more likely to be found on the edge of the microtissue showing the proliferation in the core had decreased.

Due to the decrease in cell proliferation of the HCC1806 by day 10, cell death was assessed by the TUNEL and activated cleaved-Caspase 3 staining, and in both cases a minimal number of apoptotic cells were observed in the microtissues from day 3 to day 21 (Fig. S4) which confirmed the live/dead staining. This suggests that the HCC1806 cells in the microtissues are most likely decreasing their proliferation rate, rather than undergoing apoptosis at later time points (day 10+) in the microtissues.

We speculated that the observed decrease in cell proliferation within the microtissues could indicate the establishment of oxygen and nutrient gradients within the microtissue, and potentially a hypoxic core. This has been seen in other microtissue model systems^32^. The presence of hypoxia was evaluated by the hypoxiTRAK dye, which irreversibly accumulates in cells that have experienced a hypoxic event. As shown in Figure 3, several hypoxic cells were observed starting at day 10, and increased in number till day 21. Overall, it can be concluded that the microtissue architecture changes over time. At early time points (prior to day 10) there is a proliferation of the cancer cells within the microtissue and as the cell density in the microtissue increases the proliferation slows and there is a hypoxic core formation within the microtissue.

**Figure 3.**
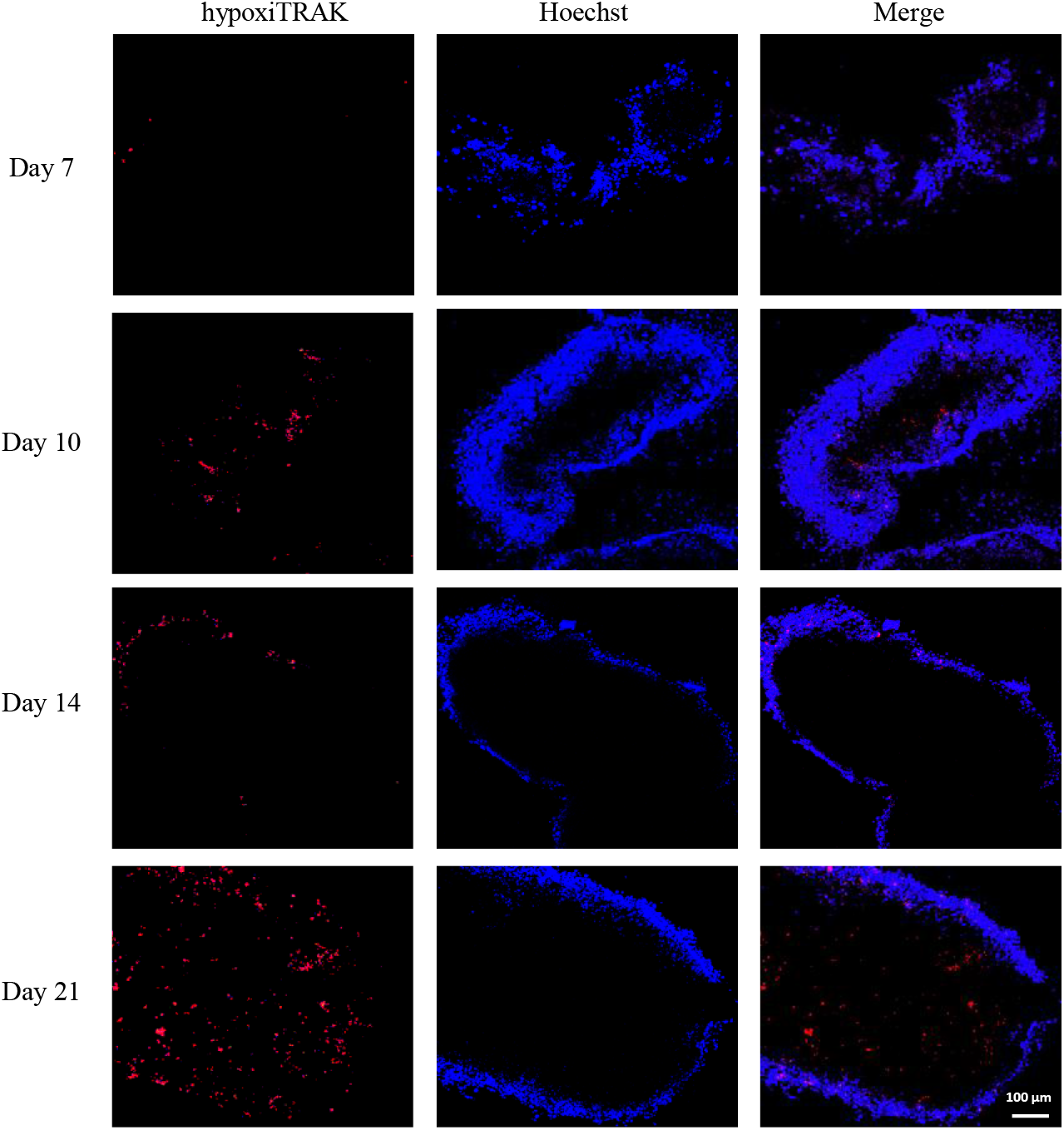
Evaluation of hypoxia. Hypoxia staining of HCC1806 microtissues at day 7, 10, 14 and 21 by the HypoxiTRAK probe (red). Microtissue were incubated with 50 nM hypoxiTRAK dye in culture media since day 5 as explained in materials and method (N=3). Nuclei were stained with Hoechst 33,342 (Blue) for 30 min before imaging. (10X)

### Cancer microtissues as a drug screening platform

An important aspect of cancer microtissue development is their utility as a drug screening platform. To evaluate their use as a drug screening platform microtissues containing either HCC1806 or MDA-MB-231 cells were screened for their response to chemotherapy drugs. Based on the H&E and proliferation data (Fig. 1 & 2) day 5 was chosen as the time point to start the drug assays. This is because the microtissues show a tissue architecture with the formation of multi-cellular groups of cells, and there is a large window of cell proliferation, and most cancer therapies target actively proliferating cells.

The response of the HCC1806 microtissues was first tested with two common chemotherapy agents, paclitaxel and doxorubicin (Fig. 4). According to the results, 2D-cultured cells were more susceptible to the tested drugs, compared to microtissue-cultured ones. Despite the fact that chemotherapeutics most often are able to completely eradicate 2D-cultured cancer cells in vitro, this does not correlate with their activity against actual patient tumors in the clinics. As shown in Figure 4, treatment of either doxorubicin or paclitaxel at the maximal plasma concentrations of the drugs^33,34^ would not have been able to completely eradicate microtissue-cultured cells.

The potency of all drugs was reduced in killing cells grown in microtissues. Additionally, efficacy was also diminished in all cases except the doxorubicin treated HCC1806 microtissues. The decline of efficacy was the highest in the case of paclitaxel, where 50% or more of the cells survived paclitaxel concentrations that were 100X the IC50, indicating the emergence of a drug resistant cell population. This resistance is not likely to be due to diffusion limitations of the drugs as doxorubicin has completely penetrated the microtissue by 10 hrs (Fig. S5) and the drug treatment duration is 4 days. Therefore, the drug resistance is more likely due to changes in the cellular phenotype that resulted from the growth in the microtissue.

Paclitaxel is an anti-tubulin agent of Taxane class, which stabilizes the microtubules by blocking disassembly of tubulins^35^. The specificity of the observed resistance to paclitaxel was tested with additional drug treatments. Starting with docetaxel, another taxane drug but with higher affinity for tubulins compared to paclitaxel^36^, it showed significantly higher potency in killing microtissue-cultured HCC1806 cells as expected. Furthermore, a different class of anti-tubulin agents, the vinca alkaloids, which target polymerization of tubulins, was tested. As shown in figure 4, neither vincristine nor vinblastine were able to eradicate all of the cells in microtissues, but the loss of efficacy of vinca alkaloids against microtissue-cultured cells was less prominent compared to paclitaxel treatment. In contrast to the basal triple negative cell line HCC1806, the mesenchymal MDA-MB-231, showed significantly higher resistance to all tested treatments (Fig. 4B). This was in accordance with the lower proliferation of the MDA-MB-231 cells and previous studies showing the dramatic difference between the sensitivity of these two cell lines to chemotherapy^37,38^. Overall, our data suggests the emergence of a paclitaxel-specific mechanism of resistance in microtissue-cultured HCC1806 cells.

**Figure 4.**
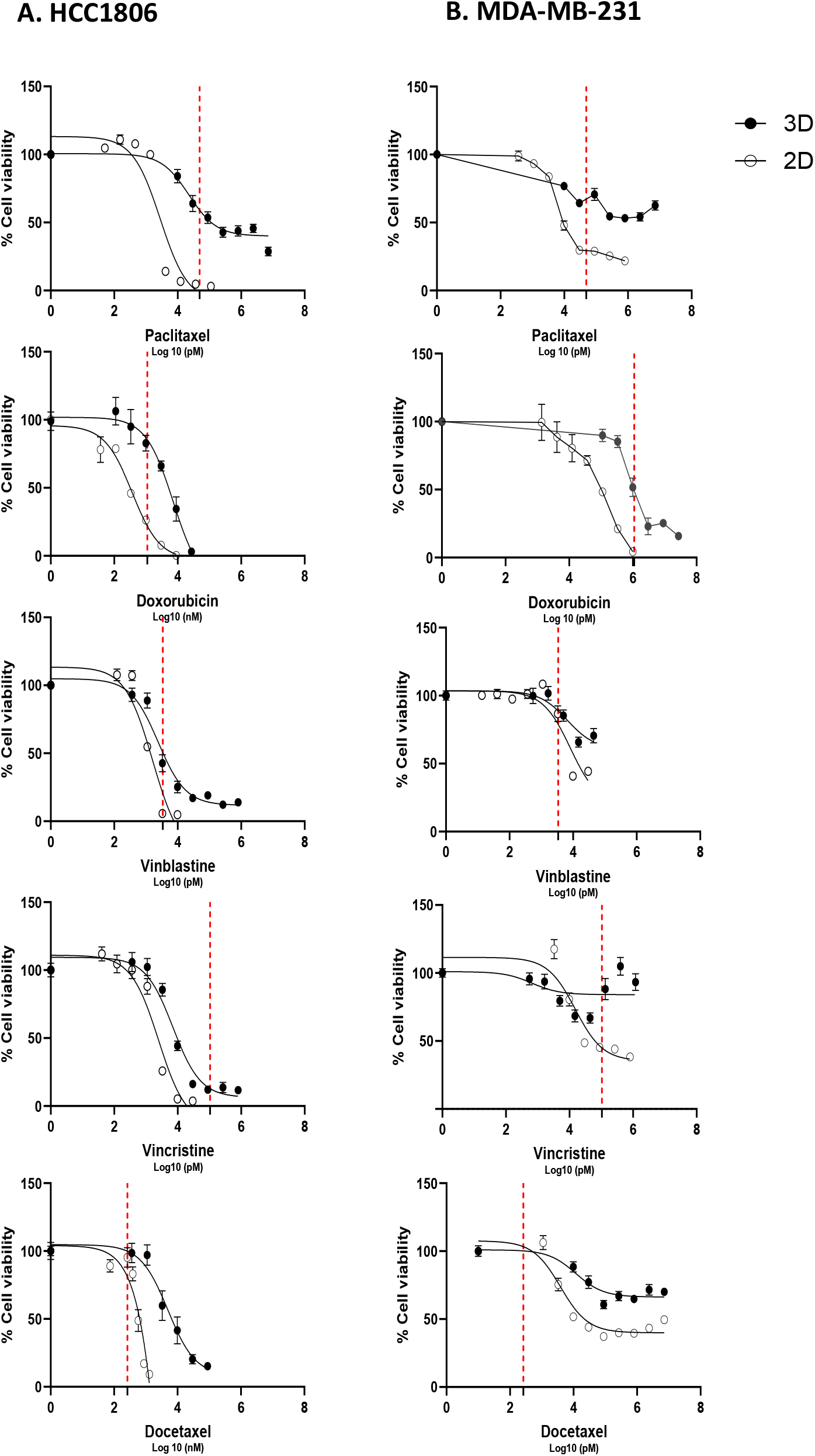
Dose-response curves of 2D-cultured vs 3D microtissue-cultured TNBC cells. HCC1806 (A) and MDA-MB-231 (B) cells were treated for 4 days with Paclitaxel, Docetaxel, Vinblastine, Vincristine and Doxorubicin. Cells were seeded in a 96-well plate at a density of 1200 cells per well for 2D-culture, and 3 microtissues per well for 3D culture. Each experiment involved three wells per condition. Data represents mean ± SEM of at least 3 experimental replicates for 3D culture, and 3 experimental replicates for 2D culture. The red dotted line represents the maximum plasma concentration of each drug which is allowed in the clinics according to the literature^33,34,43–45.^ .

## Discussion

There is no doubt that pre-clinical studies play a pivotal role in determining the success of novel drug candidates in human studies. The major biomimetic tools used in pre-clinical drug studies are simple 2D-cell culture, and animal models. The absence of complexity of human pathologies, especially cancer, in 2D models coupled with lack of human origin in animal models have created a gap in preclinical studies leading to the failure of numerous drugs in clinical trials. 3D-cell culture can be considered as a solution to better recapitulate the complexity of actual human pathology in a model with human origin. In this study, we evaluated the potentials of modular tissue engineering as a well-established technique in tissue transplantation to be used to build tumor models for cancer research. Our initial tumor model used cancer cell lines and collagen as the extracellular matrix. It could recapitulate fundamental features of tumors, such as the development of hypoxia and treatment resistance. Studying hypoxia by most of the current in vitro cancer models is based on artificial induction by using a hypoxia chamber^40,41^, where all of the cells are exposed to low oxygen levels. Conversely, our model closely mirrors natural emergence of hypoxia within an actual tumor which is derived from cell proliferation.

It is well-known that cells cultured in 2D are not able to recapitulate the therapy resistance of cancer cells of an actual patient`s tumor ^42^. As shown in figure 4, our drug treatment assays also confirmed that 2D-cultured cells are sensitive, and the applied treatments mostly led to their complete eradication. Conversely, the cells cultured in our 3D model were not responsive to the drug concentrations which were lethal for 2D-cultured cells. The emerging treatment resistance in microtissue-cultured cells led to diminished potency of all tested drugs, and in some cases, such as paclitaxel, complete loss of efficacy. Furthermore, the cancer cells cultured in microtissues recapitulated the clinical resistance of the used chemotherapeutics as the cells were resistant even at concentrations similar to the maximum plasma concentration of these drugs that is allowed in the clinics^33,34,43–45^. Therefore, this model may allow us to study resistance mechanisms to actual drug concentrations in vitro.

Previous research using modular tissue engineering has shown its flexibility to characterize an optimal microenvironment for the growth and proliferation of various cell types, including cardiomyocytes^30^, adipocytes^46^, and pancreatic islet cells^31^. Additionally, the collagen hydrogel component as the ECM, can be enriched with additional ECM components such as fibronectin and laminin^47^, based on the requirement of the tissue under study. Therefore, we will be able to improve the complexity of this model to mimic additional features of an actual tumor microenvironment. This will involve addition of non-cancerous cells, such as endothelial cells, fibroblasts, and macrophages, to mimic the stromal component and immune cell surveillance as the missing components of TME in the current model. Overall, our preliminary data have shown the capacity of this model and modular tissue engineering to fabricate a much more advanced biomimetic model of cancer but with similar ease and cost-efficiency compared to 2D-cell culture, indicating its potential to become a useful in vitro model in cancer research.

## Acknowledgements

We thank the histology core of University of Saskatchewan for their help with preparation of histology samples. This work was supported by a Saskatchewan Cancer Agency Operating grant with funds donated to the Cancer Foundation of Saskatchewan as well as a Saskatchewan Health Research Foundation (SHRF) Establishment Grant.

## Conflict of interest

All authors have declared no conflict of interests.

## Contributions

Conceptualization, M.D.C.; Research Performance, N.D., M.B., R.G., T.D., M.J.; Data analysis, N.D., M.D.C.; Writing—original draft, N.D., M.D.C.; Figures, N.D., M.D.C.; Writing—review and editing, all authors; funding acquisition, M.D.C.

## Notes

### Competing Interest Statement

The authors have declared no competing interest.

## References

1. Unger, C. et al. Modeling human carcinomas: Physiologically relevant 3D models to improve anticancer drug development. Advanced Drug Delivery Reviews 79–80, 50–67 (2014).

2. van Rijt, A., Stefanek, E. & Valente, K. Preclinical Testing Techniques: Paving the Way for New Oncology Screening Approaches. Cancers (Basel) 15, 4466 (2023).

3. Wong, C. H., Siah, K. W. & Lo, A. W. Estimation of clinical trial success rates and related parameters. Biostatistics 20, 273–286 (2019).

4. Kapałczyńska, M. et al. 2D and 3D cell cultures – a comparison of different types of cancer cell cultures. Arch Med Sci 14, 910–919 (2018).

5. Popova, N. V. & Jücker, M. The Functional Role of Extracellular Matrix Proteins in Cancer. Cancers (Basel) 14, 238 (2022).

6. Anderson, N. M. & Simon, M. C. Tumor Microenvironment. Curr Biol 30, R921–R925 (2020).

7. Wu, P. et al. Adaptive Mechanisms of Tumor Therapy Resistance Driven by Tumor Microenvironment. Front Cell Dev Biol 9, 641469 (2021).

8. Mitola, G., Falvo, P. & Bertolini, F. New Insight to Overcome Tumor Resistance: An Overview from Cellular to Clinical Therapies. Life (Basel) 11, 1131 (2021).

9. Baghban, R. et al. Tumor microenvironment complexity and therapeutic implications at a glance. Cell Communication and Signaling 18, 59 (2020).

10. Son, W.-C. & Gopinath, C. Early occurrence of spontaneous tumors in CD-1 mice and Sprague-Dawley rats. Toxicol Pathol 32, 371–374 (2004).

11. Jubelin, C. et al. Three-dimensional in vitro culture models in oncology research. Cell & Bioscience 12, 155 (2022).

12. Yakavets, I. et al. Advanced co-culture 3D breast cancer model for investigation of fibrosis induced by external stimuli: optimization study. Sci Rep 10, 21273 (2020).

13. Rodenhizer, D., Cojocari, D., Wouters, B. G. & McGuigan, A. P. Development of TRACER: tissue roll for analysis of cellular environment and response. Biofabrication 8, 045008 (2016).

14. Prince, E. et al. Microfluidic Arrays of Breast Tumor Spheroids for Drug Screening and Personalized Cancer Therapies. Adv Healthc Mater 11, e2101085 (2022).

15. Sigdel, I. et al. Biomimetic Microfluidic Platforms for the Assessment of Breast Cancer Metastasis. Front Bioeng Biotechnol 9, 633671 (2021).

16. Pape, J., Emberton, M. & Cheema, U. 3D Cancer Models: The Need for a Complex Stroma, Compartmentalization and Stiffness. Frontiers in Bioengineering and Biotechnology 9, (2021).

17. Drost, J. & Clevers, H. Organoids in cancer research. Nat Rev Cancer 18, 407–418 (2018).

18. James-Bhasin, M., Siegel, P. M. & Nazhat, S. N. A Three-Dimensional Dense Collagen Hydrogel to Model Cancer Cell/Osteoblast Interactions. Journal of Functional Biomaterials 9, 72 (2018).

19. Antoine, E. E., Vlachos, P. P. & Rylander, M. N. Review of Collagen I Hydrogels for Bioengineered Tissue Microenvironments: Characterization of Mechanics, Structure, and Transport. Tissue Eng Part B Rev 20, 683–696 (2014).

20. Szot, C. S., Buchanan, C. F., Freeman, J. W. & Rylander, M. N. 3D in vitro bioengineered tumors based on collagen I hydrogels. Biomaterials 32, 7905–7912 (2011).

21. Szot, C. S., Buchanan, C. F., Rylander, M. N. & Freeman, J. W. Cancer cells cultured within collagen I hydrogels exhibit an in vivo solid tumor phenotype. in 2011 IEEE 37th Annual Northeast Bioengineering Conference (NEBEC) 1–2 (2011). doi:10.1109/NEBC.2011.5778721.

22. Massey, A. et al. Mechanical properties of human tumour tissues and their implications for cancer development. Nat Rev Phys 6, 269–282 (2024).

23. McGuigan, A. P. & Sefton, M. V. Vascularized organoid engineered by modular assembly enables blood perfusion. Proc Natl Acad Sci U S A 103, 11461–11466 (2006).

24. McGuigan, A. P. & Sefton, M. V. Design criteria for a modular tissue-engineered construct. Tissue Eng 13, 1079–1089 (2007).

25. McGuigan, A. P. & Sefton, M. V. Modular tissue engineering: fabrication of a gelatin-based construct. J Tissue Eng Regen Med 1, 136–145 (2007).

26. McGuigan, A. P. & Sefton, M. V. Design and fabrication of sub-mm-sized modules containing encapsulated cells for modular tissue engineering. Tissue Eng 13, 1069–1078 (2007).

27. McGuigan, A. P., Leung, B. & Sefton, M. V. Fabrication of cell-containing gel modules to assemble modular tissue-engineered constructs [corrected]. Nat Protoc 1, 2963–2969 (2006).

28. Chamberlain, M. D. et al. Fabrication of micro-tissues using modules of collagen gel containing cells. J Vis Exp 2177 (2010) doi:10.3791/2177.

29. González-Callejo, P. et al. 3D bioprinted breast tumor-stroma models for pre-clinical drug testing. Materials Today Bio 23, 100826 (2023).

30. Leung, B. M. & Sefton, M. V. A Modular Approach to Cardiac Tissue Engineering. Tissue Eng Part A 16, 3207–3218 (2010).

31. Vlahos, A. E., Cober, N. & Sefton, M. V. Modular tissue engineering for the vascularization of subcutaneously transplanted pancreatic islets. Proceedings of the National Academy of Sciences 114, 9337–9342 (2017).

32. Corstorphine, L. & Sefton, M. V. Effectiveness factor and diffusion limitations in collagen gel modules containing HepG2 cells. J Tissue Eng Regen Med 5, 119–129 (2011).

33. Hertz, D. L. et al. Paclitaxel Plasma Concentration after the First Infusion Predicts Treatment-Limiting Peripheral Neuropathy. Clin Cancer Res 24, 3602–3610 (2018).

34. Harahap, Y., Ardiningsih, P., Corintias Winarti, A. & Purwanto, D. J. Analysis of the Doxorubicin and Doxorubicinol in the Plasma of Breast Cancer Patients for Monitoring the Toxicity of Doxorubicin. Drug Des Devel Ther 14, 3469–3475 (2020).

35. Weaver, B. A. How Taxol/paclitaxel kills cancer cells. Mol Biol Cell 25, 2677–2681 (2014).

36. Verweij, J., Clavel, M. & Chevalier, B. Paclitaxel (Taxol) and docetaxel (Taxotere): not simply two of a kind. Ann Oncol 5, 495–505 (1994).

37. Volk-Draper, L. D., Rajput, S., Hall, K. L., Wilber, A. & Ran, S. Novel Model for Basaloid Triple-negative Breast Cancer: Behavior In Vivo and Response to Therapy. Neoplasia (New York, N.Y.) 14, 926 (2012).

38. Abdel-Mohsen, M. A. et al. Influence of copper(I) nicotinate complex on the Notch1 signaling pathway in triple negative breast cancer cell lines. Sci Rep 14, 2522 (2024).

39. Krause, W. Resistance to anti-tubulin agents: From vinca alkaloids to epothilones. Cancer Drug Resist 2, 82–106 (2019).

40. Svanström, A. et al. The Effect of Hypoxic and Normoxic Culturing Conditions in Different Breast Cancer 3D Model Systems. Front Bioeng Biotechnol 9, 711977 (2021).

41. Ma, S. et al. Hypoxia induces HIF1α-dependent epigenetic vulnerability in triple negative breast cancer to confer immune effector dysfunction and resistance to anti-PD-1 immunotherapy. Nat Commun 13, 4118 (2022).

42. Differences in drug sensitivity between two-dimensional and three-dimensional culture systems in triple-negative breast cancer cell lines. Biochemical and Biophysical Research Communications 533, 268–274 (2020).

43. Yang, F. et al. Pharmacokinetic Behavior of Vincristine and Safety Following Intravenous Administration of Vincristine Sulfate Liposome Injection in Chinese Patients With Malignant Lymphoma. Front. Pharmacol. 9, (2018).

44. Links, M. et al. Vinblastine pharmacokinetics in patients with non-small cell lung cancer given cisplatin. Cancer Invest 17, 479–485 (1999).

45. Vermunt, M., Marchetti, S. & Beijnen, J. Pharmacokinetics and Toxicities of Oral Docetaxel Formulations Co-Administered with Ritonavir in Phase I Trials. Clin Pharmacol 13, 21–32 (2021).

46. Butler, M. J. & Sefton, M. V. Cotransplantation of Adipose-Derived Mesenchymal Stromal Cells and Endothelial Cells in a Modular Construct Drives Vascularization in SCID/bg Mice. Tissue Eng Part A 18, 1628–1641 (2012).

47. Cooper, T. P. & Sefton, M. V. Fibronectin coating of collagen modules increases in vivo HUVEC survival and vessel formation in SCID mice. Acta Biomater 7, 1072–1083 (2011).

