## Supplemental Figures for "A novel in vitro 3D cancer model based on modular tissue engineering approach"

### Supplementary Figure 1

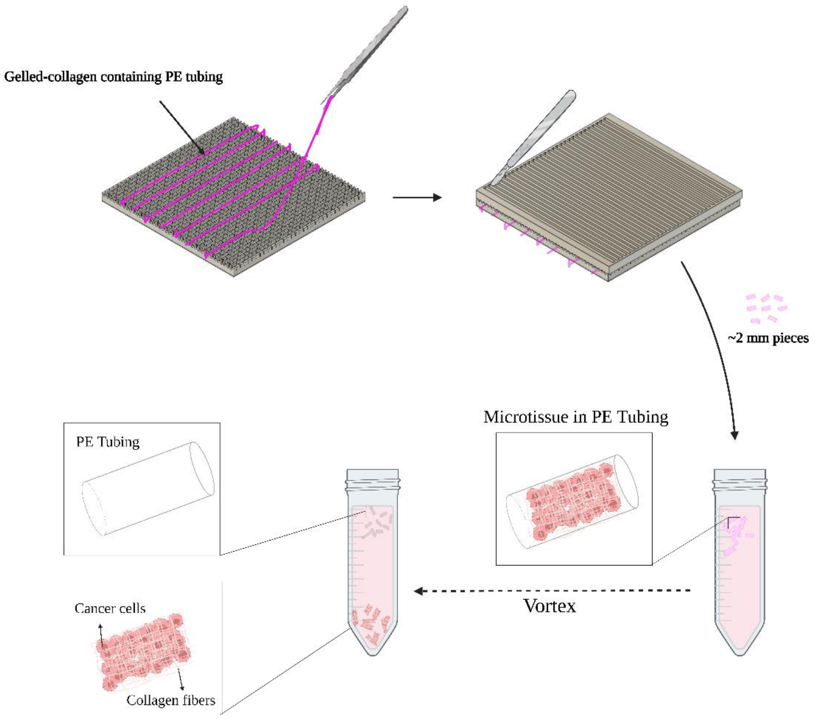


**Figure S1.** Modular tissue engineering. A mixture of collagen and cancer cells is prepared and infused into a sterile polyethylene tubing. Once collagen gel is formed, the tubing is sectioned into 2 mm modules, collected in RPMI media, and vortexed to release the microtissues from tubing. Created with BioRender.com

### Supplementary Figure 2

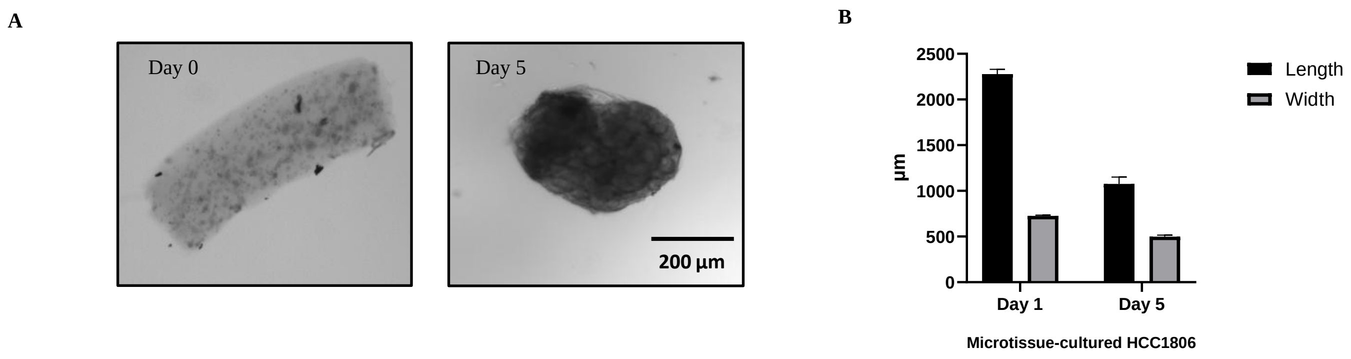


**Figure S2.** Brightfield images of a microtissue-cultured HCC1806 cells ($2\times10^6$ cell/ml of collagen) at day 1 and 5. (4X), C) Size assessment assay. The length and width of microtissues were measured by EVOS cell imager system. Each bar is a total of 45 microtissues. 15 technical replicates from 3 independent batches. Data are reported as mean ± SEM and plotted using GraphPad prism software 9.

### Supplementary Figure 3

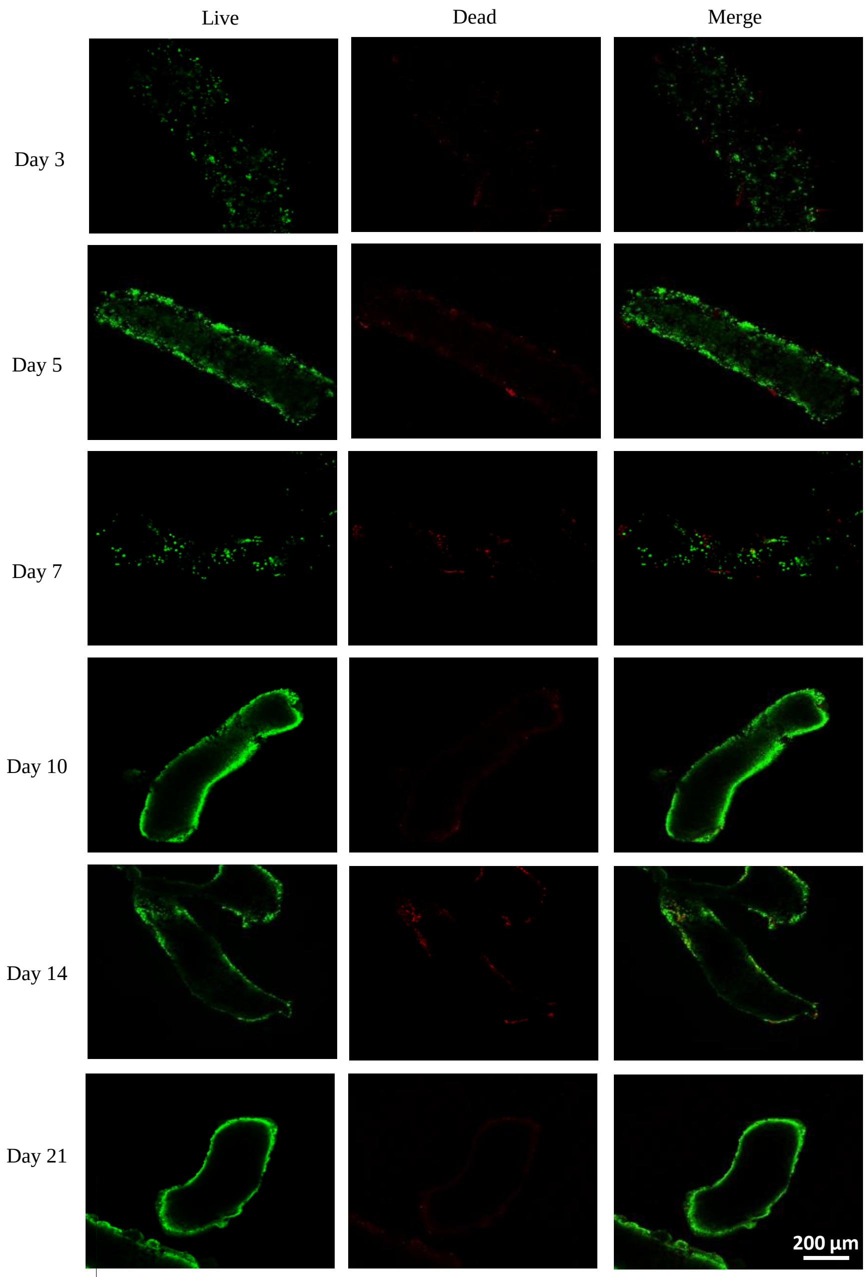


**Figure S3.** Representative images of live/dead staining of the 3D microtissue-cultured HCC1806 cells at day 3, 5, 7, 10, 14 and 21 taken by confocal microscope (N=3). The green staining is Calcein AM for live cells and the red staining is ethidium homodimer-1 for dead cells. (5X)

### Supplementary Figure 4

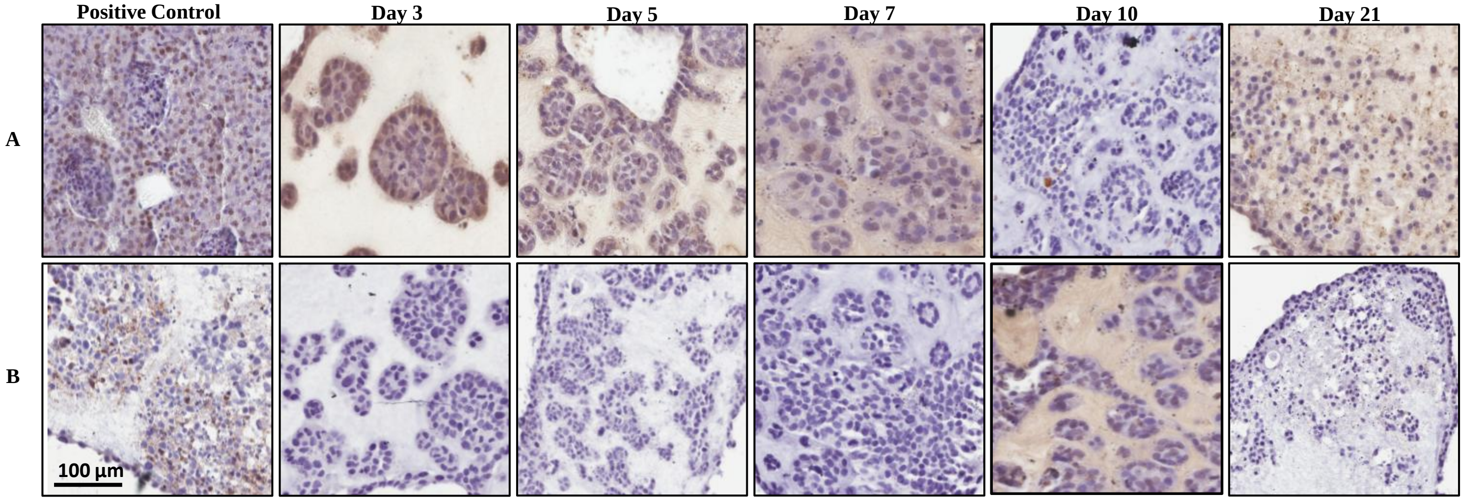


**Figure S4**. Immunohistochemical labeling of apoptotic cells in HCC1806 microtissues with Tunnel assay (A), and Cleaved-Caspase 3 antibody (B). Microtissues fixed at day 3, 5, 7, 10, and 21 were sectioned and processed with TUNNEL kit, or stained with Cleaved Caspase 3 antibody as explained in materials and methods (N=2). Mouse kidney tissue treated with DNase I, and microtissue-cultured HCC1806 treated with IC50 of doxorubicin were used as positive control in Tunnel assay and cleaved caspase 3 staining respectively. The apoptotic cells are stained brown. (20X)

### Supplementary Figure 5

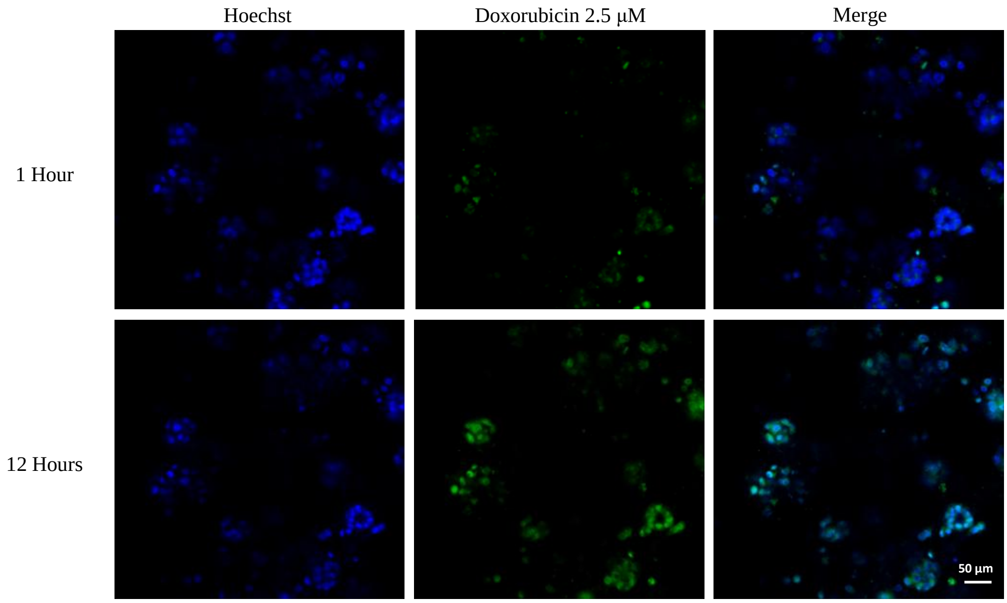


**Figure S5**. Doxorubicin diffusion assay. HCC1806 microtissues were treated with 2.5 µM concentration of doxorubicin (Green), incubated with Hoechst (Blue) for 1 hour and imaged live for 11 hours. Nuclei were stained with Hoechst 33,342 (Blue).(20X)
